# RNase H1 exists as phase-separated assemblies in association with elongating RNA Polymerase II during active transcription

**DOI:** 10.1101/2022.04.08.487625

**Authors:** Rituparna Das, Anusree Dey, Hari S. Misra, Sheetal Uppal

## Abstract

R-loops are three stranded nucleic acid structures consisting of a RNA/DNA hybrid and a single stranded displaced DNA. RNase H1 is an endonuclease which specifically degrades the RNA moiety in RNA-DNA hybrids. Here, we report that RNase H1 interacts with transcription elongation machinery during active transcription in a liquid-liquid phase separation (LLPS) dependent manner. We show that RNase H1 interacts with nascent RNA, and transcription elongation machinery in Hela cells, using in-situ nascent RNA labelling and Proximity ligation assay. Further, RNase H1 was found to exhibit properties of liquid-like condensates both in vitro and in vivo. Interestingly, RNase H1 interaction with elongating RNA Polymerase II can be disrupted by chemicals that perturb LLPS. Importantly, we show that LLPS is important for regulation of R-loop levels in the cell. Based on our results, we propose that RNase H1 exists as phase-separated assemblies in association with elongating RNA Polymerase II during active transcription.

## Introduction

R-loops are dynamic three stranded nucleic acid structures consisting of a RNA/DNA hybrid and a single stranded displaced DNA on the chromatin (Richard and Manley, 2017). Most of the R-loops are formed co-transcriptionally as confirmed by various transcription perturbation experiments (Stork et al., 2016). According to the most accepted ‘Thread back model’, R-loops are formed by invasion of the newly transcribed RNA in to the duplex DNA, displacing the coding strand (Roy et al., 2008). During transcription, R-loops may form behind transcribing RNA polymerase in case of defective or delayed mRNA processing or if the histones are slow to pack template, or by disruption of transcription-translation coupling in bacteria (Li and Manley, 2005; Domínguez et al., 2011; Wahba et al., 2011; Striling et al., 2012; Kouzminova et al., 2017). Recently, Chen et al. (2007) have demonstrated that R-loops are present in proximity to gene promoters and a free RNA end is needed for R-loop formation. Apart from transcription, RNA/DNA hybrids are also formed during replication (Burgers, 2009) and IgG Class switching (Roy et al., 2008). Presence of G rich sequences, nicked DNA or negative supercoiling favor R-loop formation (Roy and Lieber, 2009; Roy et al., 2010).

Regulation of R-loops is critical for normal cellular functioning as apart from playing important roles, undesired R-loops may pose hindrance to processes such as transcription or replication (Hamperl et al., 2017). R-loops induced by head-on collision of replication and transcription machinery, induce DNA breaks leading to genomic instability (Hamperl et al., 2017). In addition, the displaced ssDNA can be subjected to various chemical and enzymatic modifications in DNA resulting in harmful mutations (Gómez-González and Aguilera, 2007; Aguilera and García-Muse, 2012). Thus, Cells have evolved several mechanisms to combat these structures. Topoisomerases that remove negative supercoiling (Topo I) prevent R loop formation (Manzo et al., 2018). mRNA biogenesis and nuclear export factors also enable efficient RNA removal from nucleus, reducing prevalence of R loops (Li and Manley, 2005; Domínguez et al., 2011; Wahba et al., 2011; Striling et al., 2012). Helicases like senataxin, aquarius, DHX9, BLM, etc. help to resolve RNA/DNA hybrid structures in R-loops (Skourti-Stathaki et al., 2011; Chakraborty and Grosse, 2011; Sollier et al., 2014; Chang et al.,2017).

Human RNase H1 gets rid of R-loop structures by degrading RNA moiety in the R-loops by virtue of its endonuclease activity (Wahba et al., 2011; Amon and Koshland, 2016). Amongst the variety of mechanisms which cells have evolved for resolving R-loops, RNase H1 remains the primary factor which responds to high load of R-loops in a cell-cycle independent manner (Amon and Koshland, 2016). Though R-loops are known to form co-transcriptionally, the exact mechanism by which RNase H1 responds to high levels of abnormal R loops still remains to be answered. Here, we report that RNase H1 interacts with transcription elongation machinery during active transcription. We found that RNase H1 exists as liquid-like condensates in the cells. Treatment with chemicals that disrupt liquid-liquid phase separation (LLPS) leads to perturbation of RNase H1 and RNA pol II interaction as well as results in increased R-loop level in the cells. Based on these results, we propose that RNase H1 exists as phase-separated assemblies in association with elongating RNA Polymerase II during active transcription.

## Results and discussion

### Detection of in situ protein interactions with nascent RNA in a single cell using proximity ligation assay (IPNR-PLA)

While R-loops are reported to form by nascent RNA during transcription (Chen et al., 2017), visual in-situ co-localization of R-loops in the vicinity of nascent RNA has not been shown yet. We developed a highly sensitive technique for detection of in situ protein (or biomolecule) interactions with nascent RNA in a single cell using proximity ligation assay (IPNR-PLA) (Fig.1A). PLA is a assay which provides information about the proximity of two proteins in vivo using specific primary antibodies against the bio-molecules of interest giving rise to a location specific punctate fluorescent signal only if the two bio-molecules are present within 40 nm close proximity (Gullberg et al., 2004). In IPNR-PLA, Nascent RNA labeling with 5-Flourouridine (FU) followed by PLA using combination of anti-BRDU antibody and antibody against bio-moiety of interest can be used to probe bio-moiety interaction with nascent RNA. First, we labelled cellular nascent RNA by 5’-flouro-uridine (FU) during cell growth followed by immunostaining using anti-BRDU antibody. Good amount of signal was obtained indicating efficient labeling of nascent transcripts (Fig. S1A). Subsequently, we set up two IPNR-PLA reactions in Hela cells to show known interaction of nascent RNA with RNA polymerase II (RPII) and with FUS, an RNA binding protein which is known to interact with the CTD and regulate transcription elongation. FUS has been shown to regulate CTD phosphorylation during transcription elongation and help in RNA pol II pause release (Schwartz et al., 2012). Single antibody controls of anti-RPII, anti-FUS, and anti-BRDU (with FU incorporation) were also used as negative controls. The IPNR-PLA technique showed red punctate signal in the nucleus for both the reactions while all the single antibody controls did not show any signal (Fig.1B and Fig. S1B) demonstrating a strong interaction of FUS and RPII with nascent RNA, thereby validating the technique.

### R-loops co-localize with nascent transcript and RNA pol II CTD

Subsequently, we used IPNR-PLA to probe if R-loops exist in close proximity to nascent RNA in Hela cells. It is reported that D10R, E48R mutant of *E. coli* RNase H_EC_ is a catalytically dead mutant, which binds to RNA-DNA hybrid in R-loops, but does not degrade RNA, therefore, can be used as an R-loop reporter for localizing R-loops in the cell (Britton et al., 2014). Similarly, catalytically dead human Mutant RNase H1 has been used as R-loop reporter in other studies (Crossley et al., 2021; Chen et al., 2017). We used the catalytically dead *E. coli* mut-RNaseH_EC_ as an R-loop reporter in fixed and permeabilized cells to localize R-loops in IPNR-PLA as a substitute for R-loop antibody. It may be noted that this is *E. coli* mutant RNase H protein not human RNaseH1 and its purpose is to help localize R-loops in the fixed cells. The mutant protein is tagged with flag at the N-terminal end for enabling immunolabelling with anti-flag antibody for carrying out PLA. We confirmed RNA-DNA hybrid binding ability of purified mut-RNase H_EC_ by EMSA using RNA-DNA (Dig-RD) hybrid made with 5’-dig-labelled DNA (Fig. S1C). A shift in the mobility of Dig-RD hybrid confirmed that the mut-RNaseH_EC_ binds to Dig-RD hybrid without degrading the RNA moiety. Subsequently, for probing co-localization of R-loops with nascent RNA, we performed IPNR-PLA using combination of flag-tagged-mut-RNase H_EC_, anti-flag antibody and anti-BRDU antibody in FU treated cells. IPNR-PLA showed a positive fluorescent punctate signal specifically in the nucleus while the single anti-flag antibody control (anti-flag antibody with mut-RNase H_EC_) did not show any PLA signal (Fig.1B and Fig. S1D) confirming that R-loops exist in proximity to the nascent RNA in the nucleus.

Chip-seq study has shown that R-loops are present at gene promoter during transcription (Chen et al., 2017), however, in vivo co-localization of R-loops in the vicinity of transcription machinery has not been shown yet. Having confirmed R-loops proximity to nascent RNA, we wanted to check the proximity of R-loops with transcription machinery using PLA. We found significant PLA signal for colocalization of both RNA Pol II (RP II), and FUS with R-loops (Fig.1C), placing R-loops in the close proximity to the transcription machinery. We also showed known interaction of FUS with RPII (Fig.S1E). These results demonstrate that R-loops are present in the close vicinity of nascent RNA and transcription machinery.

**Figure 1.**
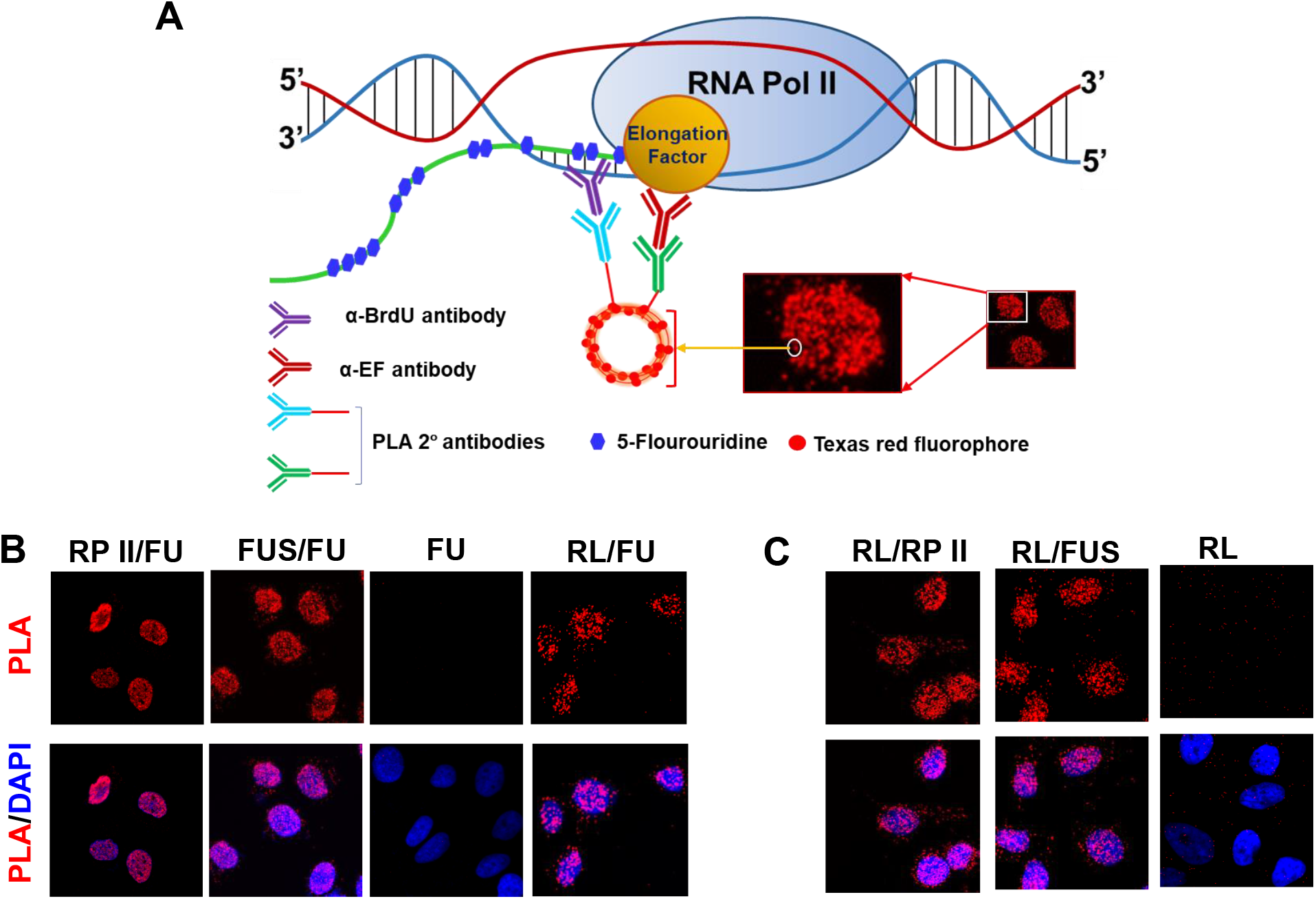
R-loops co-localize with nascent transcript and RPII in Hela cells. A. A schematic diagram showing principle of IPNR-PLA; B. IPNR-PLA showing RPII, FUS and R-loop (RL) (mut-RNase H_EC;_/anti-flag antibody) proximity to nascent RNA (FU labelling/anti-BRDU); single antibody control (FU labelling/anti-BRDU) is included as a negative control; C. PLA showing in situ association of R-loops with RPII and FUS; single antibody control R-loop (RL) (mut-RNase H_EC_/anti-flag antibody) is included as a negative control.

### RNase H1 interacts with transcription elongation machinery

RNase H1 (RNH1) is known to regulate cellular R-loops levels by degrading the RNA moiety in the RNA-DNA hybrid in R-loops (Wahba et al., 2011). Since R-loops are associated with transcription machinery, we asked if RNH1 associates with components of Transcription machinery including FUS, and RNA pol II. First, we probed RNH1 co-localization with R-loops using PLA and found low signal (Fig.2A) which might be because the catalytically active RNH1, if in close proximity, would degrade R-loops. Next, we found significant co-localization of RNH1 with RPII and FUS using PLA (Fig.2A). A negative control between RNH1 and GAPDH as well as anti-RNH1 single antibody reactions did not show any PLA signal (Fig.2A) thereby confirming that the above results show a specific association. Having demonstrated that RNH1 associates with RPII, we sought to validate and extend this analysis by determining whether RNH1 directly interacts with RPII. For this, we carried out pull down assays using recombinant proteins. Negative control having only GST beads with GST protein and flag-tagged human RNH1 did not pull down RNH1 suggesting that flag-RNH1 doesn’t bind non-specifically to the beads. Recombinant rGST-CTD (GST tagged C-terminal domain of the largest subunit of RNA Pol II) showed efficient pull down of flag-tagged human RNH1 indicating that RNH1 directly interacts with RPII CTD (Fig.2B). Thus, RNH1 interacts with the transcription machinery.

R-loops are formed due to invasion of nascent RNA in to complementary dsDNA during transcription. However, it is not known if RNH1 degrades these R-loops during transcription or post-transcriptionally. While our above results indicate the interaction of RNH1 with components of transcription machinery, it does not tell us if this interaction takes place during transcription. We found significant association of RNH1 with nascent RNA indicating at a possibility of RNH1 action during transcription (Fig. 2C). In order to further confirm this, we asked if RNH1 colocalizes with RPII during transcription elongation. Elongating RP II gets phosphorylated at ser2 and ser5 of CTD during early and late transcription elongation, respectively (Ho and Shuman,1999). The PLA results probing RNH1 interaction with elongating RPII (RPII-ser5P and RPII-ser2P) showed significant colocalization signal while the single antibody controls (RPII-ser5P or RPII-ser2P) did not show any signal (Fig. S2A), indicating that RNH1 colocalizes with elongating RP II (Fig.2C).

**Figure 2.**
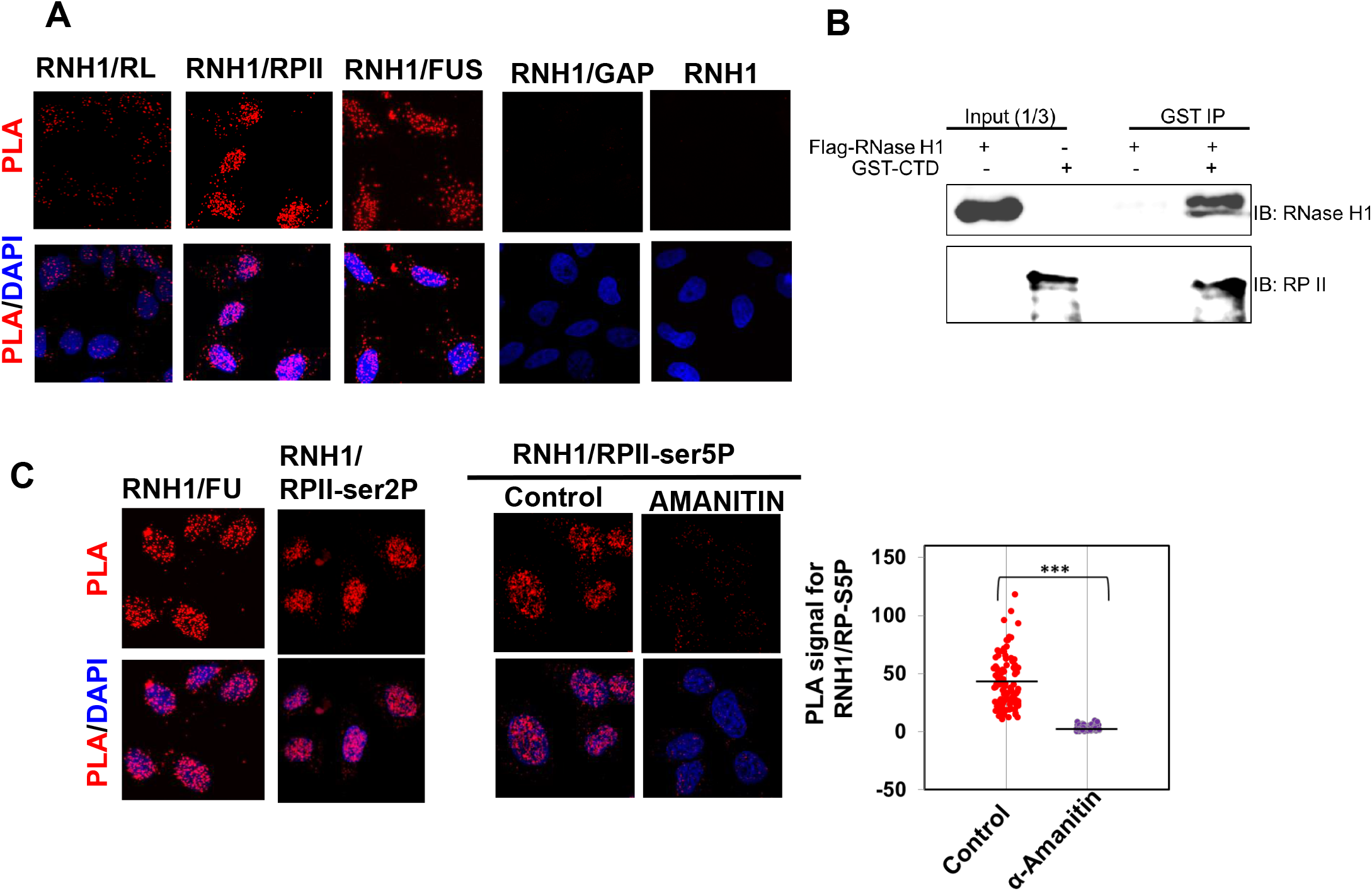
RNase H1 interacts with transcription elongation machinery. A. PLA showing association of RNH1 with R-loops, RPII and FUS; negative control for PLA using combination of anti-RNH1 and anti-GAPDH; and single anti-RNH1 antibody control is also presented; B. Immunoblots showing GST IP using recombinant GST-CTD (RPII) and recombinant flag-tagged-RNH1; C. PLA showing RNH1 interaction with nascent RNA, RPII-ser2P, and RPII-ser5P (RP-S5P) with and without α-Amanitin (50 μg/mL) treatment; corresponding stacked scatter plots with mean value as horizontal black line is presented.

Further, treatment of Hela cells with α-Amanitin, a transcription inhibitor (Fig. S2B), resulted in complete disruption of PLA signal for RNH1 with RPII-ser5P. Since ser-5 phosphorylated RPII (ser5P) is present only during transcription indicating that the PLA signal is specific in nature (Fig. 2C). Further, it is possible that RNH1 interaction with elongating RNA pol II may be mediated through co-transcriptional R-loops or nascent RNA. In order to ask this, we probed association of RNH1 with RPII-ser5P after treatment of permeabilized Hela cells with RNase A or *E. coli* RNase H_EC_ for 20 minutes before proceeding for PLA, which will result in removal of RNA or R-loops, respectively, in the cell. Both removal of RNA or RNA-DNA hybrids (R-loops) did not show any significant change in PLA signal (Fig. S2C). Another explanation for no change in the signal may be that the PLA signal is non-specific in nature and is not affected by changing conditions, however, absence of PLA signal in Fig. 2C rules out this possibility. This shows that RNH1 interaction with RPII is not mediated through R-loops. Therefore, RNase H1 interacts with elongating RNA pol II in vivo during transcription.

### RNH1 forms liquid-like condensates in vivo and in vitro

Recent advances indicate that cells utilize membrane-free RNA-protein rich condensates formed by liquid-liquid phase separation (LLPS) as compartments for organizing the intracellular space to partition diverse set of biomolecules selectively (Banani et al., 2017). Phase separation has emerged as a major driver of functional compartmentalization within the cells, allowing rapid and dynamic isolation of specific activities from the surrounding cellular environment, without the need of a membrane. One such example is formation of transcription factories, which are biomolecular condensates containing RNA polymerase, at the discrete sites where transcription occurs in the cell nucleus (Razin et al., 2011). Both FUS and CTD have disordered protein domains which are known to form macromolecular condensates via LLPS (Boehning et al., 2018; Burke et al., 2015). Interestingly, the PONDR score (Xue et al., 2010), a prediction of the degree of disorderedness, for FUS is ∼ 0.71 and for RNH1 is ∼ 0.41 while it is ∼0.23 for an unrelated protein (PCNA). The predicted three-dimensional structures (Jumper et al., 2021) of both FUS and RNH1, also reveal the disordered region (Fig. 3A). Notably, the disordered linker region in RNH1, between amino acids residues ∼70 to ∼130, connecting the two globular domains is well defined with a sharp transition from order to disorder (score > 0.5) and again back to order (Fig. 3A and Fig. S3A). The presence of the disordered region in RNH1 suggests that it may form liquid like condensates. We asked if RNH1 forms liquid like droplets in vitro. We observed purified human RNH1 under microscope in DIC mode and found that RNH1 formed droplets of various sizes by itself in the buffer (Fig. 3B and Fig. S3B). There were no droplets present in the buffer with only Sypro indicating that the droplets are from recombinant RNase H1 protein (Fig. S3B). Further, to test the dynamic nature of these droplets, we carried out Fluorescence recovery after photo-bleaching (FRAP) analysis using Sypro staining of the purified human RNH1. FRAP is a way to determine the molecular dynamics and mobility of the phase-separated liquid droplets where a small region of interest in the protein droplet is photo-bleached with a laser. The fluorescence recovery is quick in liquid-like droplets, slow in gel-like droplets, and the signal fails to recover in protein aggregates after photo-bleaching. We measured FRAP using Sypro stained RNH1 droplet and found that the fluorescence recovery was very quick (∼80 % recovery within 24 seconds) with t_1/2_ of 3.5 seconds, indicating the liquid-like nature of these droplets. (Fig. 3C and Movie S1). These results confirmed that RNH1 exists as phase separated liquid like droplets in vitro.

**Figure 3.**
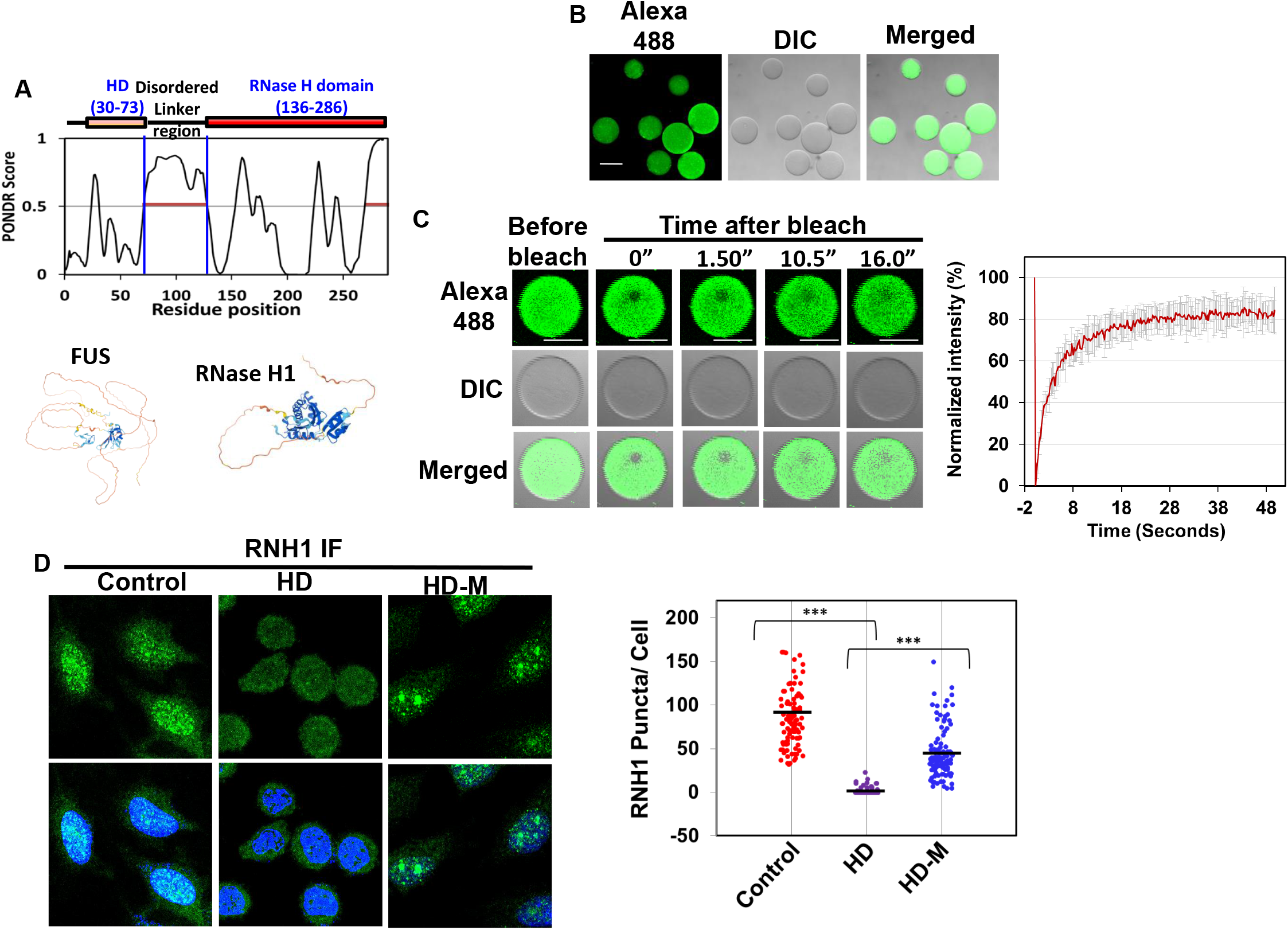
RNH1 forms liquid-like condensates in vivo and in vitro. A. PONDR (Xue et al., 2015) Plot aligned with different domains of RNH1. Predicted AlphaFold tertiary structures (Jumper et al., 2021) of RNH1 and FUS showing disordered region; B. DIC and fluorescent confocal images (40 X objective) using Sypro labelled purified RNH1 protein (also see Fig. S3B) C. FRAP assay using Sypro labelled purified RNH1 protein; Plot between average percentage recovery with standard deviation vs time (see methods for details). Scale bar (50 µm) is shown by white line. This image is grabbed by FRAP module in confocal microscope (Olympus) with maximum allowed resolution 512 × 512 µm, therefore, the edges look rough because of poor resolution. D. RNH1 immuno-staining in untreated (control), 1,6-hexanediol (HD) treated, HD followed by media addition (HD+M); corresponding stacked scatter plots with mean value as horizontal black line is presented; Number of Puncta were quantified using ImageJ (described in details in material and methods) (Schneider et al., 2012), also see Fig. S3C.

To confirm if RNase H1 exists as liquid condensates in vivo, immunofluorescence analysis was carried out. Here, RNH1 exhibited puncta staining, mainly in the nucleus (Fig. 3D and Fig. S3C). Further, a good tool to probe the physical properties of membrane-less compartments is treatment with 1,6-hexanediol (HD), which disrupts hydrophobic and non-ionic interactions responsible for formation of LLPS in the cell (Kroschwald et al., 2017). In order to probe the nature of these puncta, cells were treated with 1,6-hexanediol (HD) for short duration, which is not likely to disrupt the cellular physiology (Kroschwald et al., 2017) and stained for RNase H1 immunofluorescence. Interestingly, there was a significant reduction in the number of nuclear puncta staining upon HD treatment (Fig. 3D) indicating that RNase H1 may exist as liquid-liquid phase separated condensates in the nucleus. Next, in order to test if the loss of puncta is reversible, we incubated the cells in complete media for one hour after HD treatment (Fig. 3D). The RNH1 nuclear puncta staining also reappeared after washing with excess of complete media, indicating that RNH1 exists in form of dynamic liquid-like condensates in the cell. These results confirmed that RNH1 exists as phase separated liquid like condensates in vivo and in vitro.

### RNH1 exists as phase separated assemblies in association with elongating RPII

Our present results and others’ work show that both RNH1 and RPII (Boehning et al., 2018) exist as phase separated condensates in the cell and also interact with each other. Next, in order to ask if RNH1 interaction with elongating RPII is dependent on LLPS, we used PLA to test the effect of HD on the co-localization of RNH1 with RPII and RPII-ser5P. We found that, HD treatment lead to a drastic decrease in the co-localization of RNH1 with RPII (Fig.4A) as well as with RPII-ser5P suggesting that RNH1 interaction with elongating RPII might be through LLPS. In order to further confirm this, we asked if RNH1 condensates (puncta) colocalize with elongating RPII (RPII-ser5P) in the cell using immunostaining. For this, Hela cells were immunostained with RNH1 and RPII-ser5P specific primary antibodies from different species followed by secondary antibodies conjugated to different fluorophores (Fig.4B). The images from immunostained cells were assessed for the spatial overlap of the two fluorescent labels to test if elongating RPII colocalizes with RNH1 condensates, using ImageJ plugin JACoP (Bolte and Cordelieres, 2006). The Pearson’s correlation coefficients (PCC) was calculated and it was found that ∼45 % cells showed PCC above 0.8 and more than ∼75 % cells showed PCC value above 0.6, which suggests a strong correlation indicating significant colocalization of RNH1 condensates with elongating RNA pol II in the nucleus. These results indicate that RNH1 exists as phase separated assemblies in association with elongating RPII in the cell.

### LLPS is critical for regulation of R-loops in the cell

It has been reported that majority of the R-loop interactome proteins have intrinsically disordered region which may exist as LLPS foci (Dettori et al., 2021). After having confirmed that RNH1 exists as phase separated assemblies in association with RPII, we asked if there is an effect on R-loop levels upon disruption of LLPS by HD. For this, we monitored R-loop level with and without HD treatment in Hela cells, using immunofluorescence with S9.6 antibody, which binds to DNA-RNA hybrids with very high affinity. In untreated cells, there was very low nuclear S9.6 staining with cytoplasmic background signal, which is reported to be due to the presence of RNA-DNA hybrids present in mitochondria (Smolka et al., 2021). Upon HD treatment, there was significant increase in S9.6 staining both in the nucleus and in cytoplasm (Fig. 4C). In order to make sure that the observed signal is specifically due to increase in RNA-DNA hybrids (R-loops) in the cell, we treated both the control and HD treated cells with *E. coli* RNase H_EC_ after fixing and permeabilizing the cells but before antibody incubation step to specifically dissolve the RNA-DNA hybrids. RNase H_EC_ will result in depletion of any signal specifically present due to RNA-DNA hybrids. Both in control and HD treated cells, most of the cytoplasmic signal was unaffected by RNase H_EC_ treatment. In HD treated cells, RNase H_EC_ treatment showed a significant reduction in the nuclear S9.6 staining specifically (Fig. 4C), indicating that the increase in nuclear signal is due to higher level of R-loop in the cell upon HD treatment. The increase in R-loop level might also be due to direct inhibition of RNH1 catalytic activity in presence of HD. However, we did not find any change in purified RNH1 catalytic activity in presence of HD (Fig. S3D), in vitro, thereby ruling out this possibility. In the absence of a direct effect on the catalytic activity in vitro, we speculate that the increase in R-loops upon HD treatment might be due to decreased accessibility of RNH1 to R-loops under in-vivo conditions. Therefore, disruption of LLPS by HD treatment in Hela cells resulted in significant increase in nuclear R-loops level indicating that LLPS is critical for regulation of R-loops (Fig.4B).

**Figure 4.**
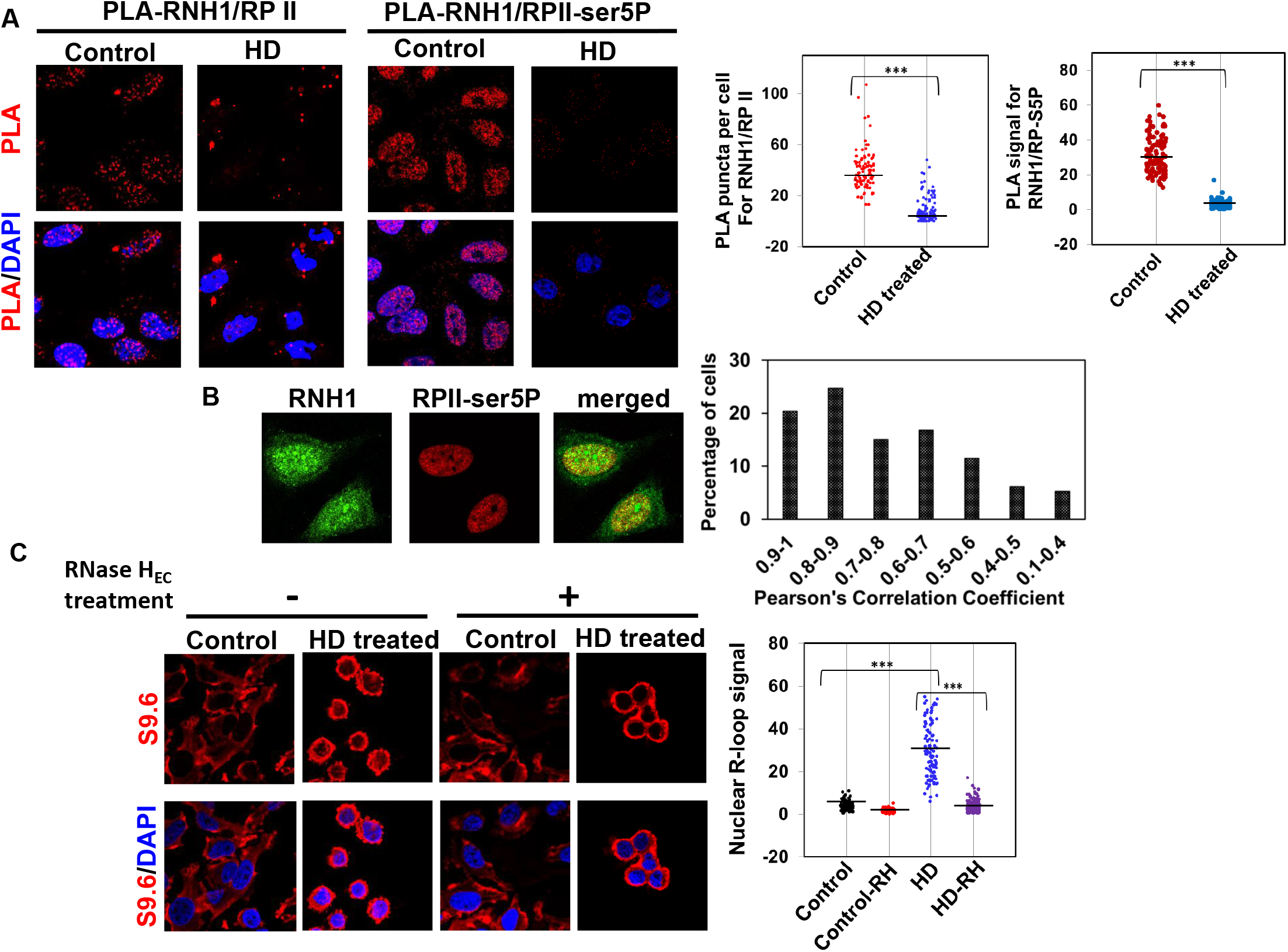
RNH1 exists as phase separated assemblies in association with elongating RPII A. PLA showing RNH1 interaction with RPII and RPII-ser5P in Hela cells, with and without HD treatment. The PLA intensity in the images was measured using ImageJ software (described in details in material and methods) (Schneider et al., 2012). B. Representative confocal images of Immunostained Hela cells showing colocalization of RNH1 with RPII-ser5P (RP-S5P). A plot showing person ‘s correlation coefficient vs percentage of cells for colocalization of RNH1 with RPII-ser5P. C. S9.6 immunostaining in cells with and without HD treatment, with and without RNase H_EC_ treatment; Corresponding stacked scatter plots with mean value as horizontal black line are presented.

Taken together, we make an important observation that RNH1 exists as dynamic liquid like condensates in the cell and its interaction with TE machinery is dependent upon liquid-liquid phase separation. R-loops are known to form in actively transcribed genes. However, presence of R-loops in the actively transcribed genes does not necessarily mean that RNH1 would interact with Transcription machinery or associate with it during active transcription. There is always a possibility of RNH1 acting post-transcriptionally. Our results show for the first time that RNH1 interacts with elongating RPII during active transcription elongation. Our results also show that RNase H1, one of the major R-loop processing enzyme, exist as liquid like condensates. In line with this, in a theoretical analysis, Dettori et al. (2021) predict that R-loop interactome proteins have intrinsically disordered region and exist as LLPS foci (Dettori et al., 2021). Further, promoter-proximal pausing results in R-loop formation (Chen et al., 2017). It is possible that RNH1 association with paused RPII (Ser5P) is necessary for resolution of these R-loops for RPII promoter clearance. R-loops also lead to de novo anti-sense transcription by acting as intrinsic pol II promoters which may have role in gene regulation (Tan-Wong et al., 2019). RNH1 overexpression has been shown to selectively down-regulate the anti-sense transcription by degrading R-loops (Tan-Wong et al., 2019). RNH1 interaction with the transcription elongation machinery might be important to regulate the levels of these R-loops and regulate anti-sense transcription. These reports (Chen et al., 2017; Tan-Wong et al., 2019) suggest an important role for RNH1 during transcription. The present study extends these findings and provides new evidence for a cross-talk between transcription and RNH1 role in the cell. Interestingly, higher R-loops levels have been observed in RNA metabolism neurological disorders associated with mutations that are known to disrupt phase separation in FUS and other IDR proteins (Salvi et al., 2015; Conicella et al., 2016). It remains to be explored if RNH1 phase separation has a role to play in these pathological conditions. Based on our results that RNH1 exists as liquid like condensates which are phase separated; RNH1 interaction with transcription elongation machinery depends on liquid-liquid phase separation; and disruption of LLPS leads to increased R-loops level in the cell, we speculate that RNH1 interaction with transcription elongation machinery might help in nucleation of RNH1 at the R-loop sites during transcription.

## Acknowledgments

We acknowledge financial support from Bhabha Atomic Research Centre, Department of Atomic Energy, Mumbai, India.

## Author contributions

RD performed experiments, analyzed data and wrote the manuscript, AD provided reagents and assisted with data analysis, HSM reviewed the manuscript and gave constructive comments, SU conceived the study, analyzed data, guided experiments and wrote the manuscript. All the authors approved the final version.

## Materials and methods

### Cell culture and plasmids

HeLa cells were cultured in DMEM medium supplemented with 10% fetal bovine serum. Primers (Table. S1) were synthesized from IDT, USA and Europhin, Germany. The bacterial expression plasmid for Flag-tagged human RNase H1, pET-11d-flagRNase H1, was generated by replacing taf7 gene fragment in pET11d-flag-taf7 (Wu and Chiang, 1996) by RNaseH1 (+1 to +784, without mitochondrial localization signal sequence) at Nde1 and BamH1 sites. RNaseH1 was PCR amplified using human RNase H1 cDNA in pENTR221 plasmid (Ultimate ORF clones, IOH4870, ThermoFisher Scientific) as a template. The *E. coli* mut-RNase H_EC_ (D10R, E48R) was PCR amplified using pICE-RNaseHI-D10R-E48R-NLS-mCherry vector as template. The bacterial expression plasmid, pET-11d-mut-RNase H_EC_ (D10R & E48R), for Flag-tagged *E. coli* mut-RNase H_EC_ (D10R, E48R) was generated by replacing taf7 gene fragment in pET11d-flag-taf7 (Wu and Chian, 1996) by mut-RNase H_EC_ at Nde1 and BamH1 sites. pICE-RNaseHI-D10R-E48R-NLS-mCherry vector (Britton et al., 2014) was a gift from Patrick Calsou (IPBS Institut de Pharmacologie et de Biologie Structurale, Toulouse, France). The human GST-CTD construct in pGEX vector was a gift from Dinah Singer (National Cancer Institute, National Institutes of Health, Bethesda, MD). All the plasmid clones created during this study were confirmed by sequencing.

### Proximity ligation assay (PLA)

PLA is a highly sensitive technique which provides information about the proximity of two proteins in vivo using specific primary antibodies against the bio-molecules in question, oligo-conjugated secondary antibodies and a series of ligation, polymerization reactions, which gives rise to a punctate fluorescent signal only if the two bio-molecules are within 40 nm close proximity (Gullberg et al., 2004). For PLA, approximately 10^4^ Hela cells were grown overnight in microchamber slides [µ-Angiogenesis Slide (I-bidi)]. PLA was conducted using the Duolink® In Situ PLA® Kit (Sigma) according to the manufacturer’s protocol. For co-localization of R-loop with other proteins, cells were incubated with 5 µg of flag-mut-RNaseH (D10R, E48R)/ well, after fixing and permeabilization for binding to R-loops (3). This was followed by PLA using combination of anti-flag antibody and primary antibody specific to protein of interest for which co-localization needs to be ascertained. For testing colocalization during transcription inhibition, cells were treated with α-Amanitin (50 µg/ml) (Bushnell et al., 2002) for 2 hours before proceeding for PLA. For 1,6-Hexanediol (HD) treatment, growing cells were incubated with 3 % HD in media for 15 minutes before proceeding for PLA. For single antibody controls, only one primary antibody was added to the PLA reaction. For RNase H (Sigma)/ RNase A treatment before PLA, growing cells were permeablized using 1.0 % tween-20 for 10 minutes and incubated with RNase H_EC_ (0.5 units/ µl) or RNase A (1 µg/µl) in media for 20 minutes before proceeding for fixation and PLA.

For observing co-localization of nascent RNA and a specific protein (IPNR-PLA), growing Hela cells were treated with 5-flourouridine (FU) (2 mM) for 30 minutes for nascent RNA labelling. Subsequently, PLA was carried out using anti-BRDU antibody and specific primary antibody against protein of interest. The primary antibodies used in PLA are as follows: rabbit polyclonal anti-Fus antibody, 1:100 (HPA008784 of Sigma), mouse monoclonal anti-Fus antibody, 1:100 (AMAb90549 of Sigma), rabbit polyclonal anti-Flag antibody,1:100 (F7425 of Sigma), mouse monoclonal anti-RNase H1, 1:100 (sc-376326 of Santa Cruz Biotechnology), mouse monoclonal anti-BrdU, 1:200 (B2531 of Sigma), rabbit BRDU, 1:100 (ab152095 of abcam), mouse monoclonal RNA Pol-II (8WG16), 1:100 (sc-56767 of Santa Cruz Biotechnology), rabbit polyclonal anti-RNase H1, 1:100 (ab229078 of abcam), rat monoclonal anti RNAP CTD beta subunit phospho ser-5, 1:100 (04-1572 of Millipore), rat monoclonal anti RNAP CTD beta subunit phospho ser-2, 1:100 (04-1571 of Millipore). Cells were imaged with Olympus Fluoview Confocal Laser Scanning Microscope, (Model FV3000) under 63X objective with immersion oil. For PLA between the following combinations RP II/FU; RNH1/RPII-ser2; and RNH1/RPII-ser5; single plane images were recorded. For remaining PLA reactions, Z-stacks were collected and maximum intensity projection images were used for presentation and intensity calculations. The PLA intensity in the images was measured using ImageJ software (Schneider et al., 2012). For signal measurement, wherever countable puncta were there, the number of Red fluorescent puncta in the nucleus were counted, however, where puncta were difficult to count due to merging, it was measured as Red fluorescence signal intensity corresponding to the nuclear region. For measuring RNase H1/RPII-ser5 PLA intensity, DAPI stained region was selected by free hand as Region of Interest (ROI) and corresponding red florescence in the ROIs was measured. For measuring PLA puncta in RNase H1/RP II sample, DAPI stained region was selected as ROIs and numbers of puncta were counted with Analyze particle function of Image J.

### Preparation of recombinant proteins

Recombinant GST-CTD (RP II) was induced in *E. coli* cells (BL21 Rosetta-gami strain; Novagen) using 0.2 mM isopropyl_β-D-1-thiogalactopyranoside (IPTG) for 4 hours at 30 °C and purified on Glutathione Sepahrose beads (GE). The protein was eluted using glutathione buffer (8 mg/ml of reduced glutathione in 50 mM Tris-HCL, 100 mM NaCl, pH: 8.0). Recombinant Flag-tagged RNase H1 or Flag-mutant-RNase H_EC_ were induced in the similar manner in *E. coli* cells using 0.2 mM IPTG for 4 hours at 30 °C and purified on flag beads (Merck) and eluted using flag peptide buffer (200 µg/ml flag peptide in 10 mM Tris-HCL, 150 mM NaCl, pH: 8.0). After elution, glutathione or flag peptide were removed by buffer exchange and the purified proteins were concentrated on Amicon Ultra 50K filters (Millipore). Purified proteins were recovered in 20 mM Hepes, pH 7.9, 100 mM NaCl, 0.2 mM EDTA, and 20 % vol/vol glycerol and stored at -80 °C.

### Electrophoretic mobility shift assays

Electrophoretic Mobility Shift Assay (EMSA) was performed with a 25 bp long RNA/DNA hybrid. The single strand DNA oligo was first labelled with Digoxigenin at 5’-end using end labelling kit (Roche) as per the manufacturers’ protocol. For making DNA-RNA hybrids, equimolar quantity of Dig-labelled ssDNA and RNA were annealed (10 minutes at 94 ^°^C followed by gradual cooling to room temperature) to form Dig-RNA/DNA (dig-RD) hybrid. After incubating 0.1 µM of dig-RD hybrid with 0.2 µM of protein in binding buffer (25 mM Tris-HCl pH 7.5, 1 mM DTT, 10 mM EDTA, 50 μg/ml BSA, 50 mM KCl) for 15 minutes, the nucleoprotein complex is resolved by electrophoresing on a 6% PAGE followed by electro-transfer to Nylon N+ membrane. The membrane was exposed to UV light for crosslinking RNA-DNA hybrid and subsequently processed for dig immunostaining giving rise to blue colour signal as per the manufacturers’ protocol using anti-DIG antibody conjugated with alkaline phosphatase and colorimetric substrate [NBT/BCIP (5-bromo-4-chloro-3-indolyl-phosphate/nitro blue tetrazolium].

### Pull down assays and Immunoblotting

For pull down assays, GST-agarose magnetic beads (Merck) were incubated with equimolar concentration of flag-tagged human RNase H1 and GST-CTD proteins in PBS buffer (pH: 7.4) for 3 hrs at 4 °C. A negative control of GST beads with GST and RNase H1 was also used to check non-specific binding of RNase H1 to beads. Following incubation and washing (50mM Tris (pH 8.0), 150 mM NaCl, and 0.2% NP-40), proteins were separated on SDS/PAGE gels and transferred onto a PVDF membrane (Merck). After blocking the membrane with 5% fat-free milk in TBST, the blots were incubated with the appropriate primary antibody. The primary antibodies used were anti-RP II mouse monoclonal antibody antibody (sc-56767 of Santacruz Biotechnology) and rabbit polyclonal anti-RNase H1 (ab229078 of abcam). The secondary antibodies IRDye®800CW or IRDye®680RD (LI-COR Biosciences) were used for protein detection. The immunoblot images were captured using the Odyssey XF Dual-Mode Imaging System (Licor, USA).

### Immunofluorescence imaging

For immunostaining with anti-BrdU (1:200, B2531 of Merck), anti-RPII-ser5P (1:1000, ab5408 of abcam) and R-loop antibody (S9.6, 1;100, MABE1095 of Millipore), Hela Cells grown on cover slips were fixed with 4% paraformaldeyhde, permeablized with 0.5% Triton-X. For nascent RNA visualization, cells were first incubated with 5-flourouridine (FU) (2mM) in media for 30 minutes before fixing, unless otherwise mentioned. For RNase H1 (1:100, ab229078 of abcam) immunostaining, cells were fixed and permeabilized using ice-cold 100% methanol followed by ice-cold acetone. This was followed by blocking and incubation with appropriate primary antibodies overnight at 4 °C and species specific secondary antibodies [polyclonal Alexa fluor 488 goat anti-rabbit,1:1000 (A11070 of Invitrogen Biosciences), polyclonal Alexa fluor 555 rabbit anti-mouse,1:1000 (A21427 of Invitrogen Biosciences)] for 1 hour at room temperature. Cells were then stained with DAPI anti-fade mounting media (Merck). For 1,6-Hexanediol (HD) treatment, growing cells were incubated with 3 % HD in media for 15 minutes before proceeding for fixation. For HD treatment followed by media, cells incubated with 3 % HD for 15 minutes were washed with excess of media and incubated in media for 1 hours followed by fixation and RNase H1 staining. For calculation of RNase H1 puncta in cells, Image J software was used. For all the RNase H1 immunostained images corresponding to control and treated samples, images were converted to 8 bit black and white images and a cut off threshold of 19-224 was set, followed by measurement of number of RNase H1 puncta in DAPI stained region of interests of individual cells using the analyze particle function. For RNase H_EC_ treatment during R-loop staining, cells were fixed, permeabilized and incubated with 0.5 units/ µl of RNase H_EC_ (M0297S of NEB) before immunostaining. Confocal single plane images were acquired using Olympus Fluoview Confocal Laser Scanning Microscope (Model FV3000) under 63X objective with immersion oil. The fluorescence intensity was measured using ImageJ software.

### Confocal Imaging of droplets of purified RNH1 and FRAP

Purified Protein (10 µM in buffer) was mixed with Sypro orange (5X), dispensed into a well of microchamber slides [µ-Angiogenesis Slide (I-bidi)] and kept for 10 minutes at room temperature for dye to label the protein. This mix was imaged using Olympus Fluoview Confocal Laser Scanning Microscope (Model FV3000) under 10X objective unless otherwise mentioned. For FRAP, a small region of interest (ROI) was selected and bleached using laser at 15 % intensity. Three ROIs were selected for FRAP analysis using manufacturer’s protocol. A third ROI was placed in an unbleached area which was used during calculations for normalization of intensity for compensation of the overall signal bleach due to laser exposure during the time lapse. Similar sized droplets of comparable intensities were chosen and bleached using the constant area of ROI and detector settings. Bleach experiments were recorded at interval of 250 ms using FRAP module of FlouView and quantified using the FRAP module of the Cell Sense software (Olympus). For each time point, the fluorescence intensities within the bleached ROI were normalized to the fluorescence intensity of the corresponding unbleached ROI. Also, for simplicity, all the values were scaled with respect to the zero time value by subtraction. Normalized intensities were then used to calculate percentage recovered fluorescence with respect to the fluorescence intensity before bleaching as base value and plotted against time. At least six individual ROIs from different cells were analyzed and plotted using Microsoft excel (mean ± SD).

### Colocalization analysis

Colocalization between RNase H1 with RPII-Ser5P in the cell nucleus was determined from the acquired images using JACoP plugin of the ImageJ analysis tool and expressed as Pearson’s correlation coefficient (PCC). At least 100 cells were taken for analysis. A plot between percentage cells and PCC is presented.

### In vitro RNase H1 catalytic activity assay

The RNA/DNA hybrid substrate was generated by mixing equimolar quantity of RNA with a complementary DNA oligo by heating at 94 ^°^C for 10 minutes followed by gradual cooling to room temperature and stored in 4 ^°^C. For determining the activity of purified human RNase H1 in vitro, 0.5 µM of the substrate was incubated with 100 nM of hRNase H1 in presence of 50 mM NaCl, 25 mM Tris-HCl pH 7.5, 5 mM MgCl_2_, 5 mM DTT, 50 μg/mL BSA with and without 5% 1,6-Hexandiol as indicated. The digested products were resolved on 10 % native PAGE at constant voltage of 150V for 20 minutes, followed by post-staining of the gel with EtBr. Image of the gel was captured under UV light using a gel documentation system (Geldoc Go imaging system, Bio-Rad).

### Statistical analysis

The stacked scatter plots were plotted using fluorescent signal or puncta number (as indicated) from scoring at least 100 cells for each sample, the p-value for statistical significance was obtained using student’s t-test (two-tailed, unpaired). All the experiments have been repeated at least thrice and representative results are shown.

## Supplemental information

### Legends to Supplementary Figures

**Fig. S1.**
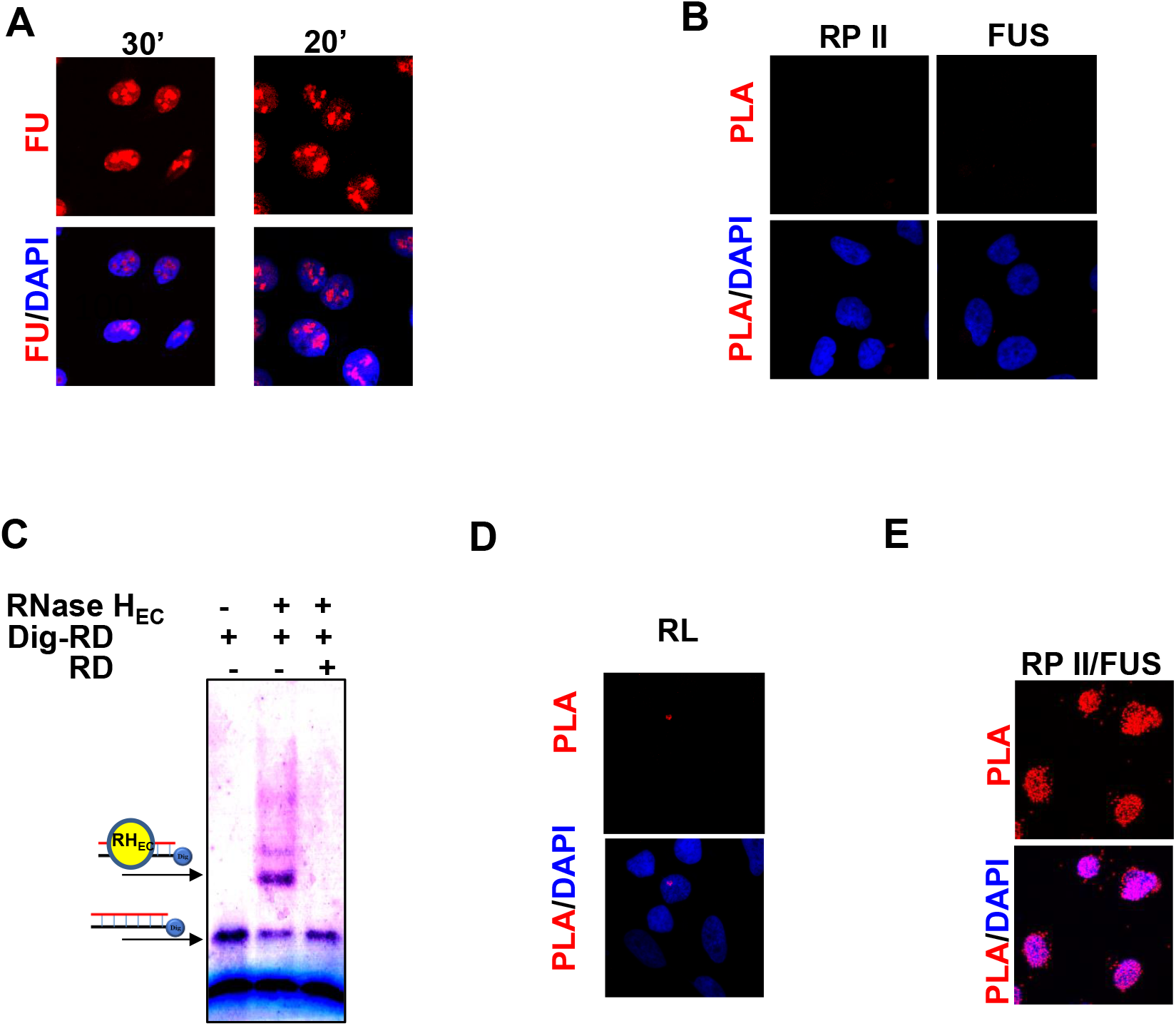
A. Nascent transcript labelling using 5’-Flourouridine (FU) in Hela cells. Growing cells were treated with 2mM FU for the indicated times and then subjected to immnunostaining with anti-BrdU antibody. PLA showing B. Single antibody controls of anti-RPII and anti-FUS; D. Single antibody control R-loop (RL) (mut-RNase H_EC_/anti-flag antibody) as negative controls; E. interaction of FUS with RPII; C. EMSA showing RNA-DNA hybrid binding ability of *E. coli* mutant (D10R, E48R) RNase H_EC_ using 5’-dig-labelled DNA-RNA (RD) hybrid. Note that excess of unlabeled RD in lane 3 leads to complete depletion of shifted band due to displacing of the Dig-RD with excess unlabeled RD indicating the specificity of binding.

**Fig. S2.**
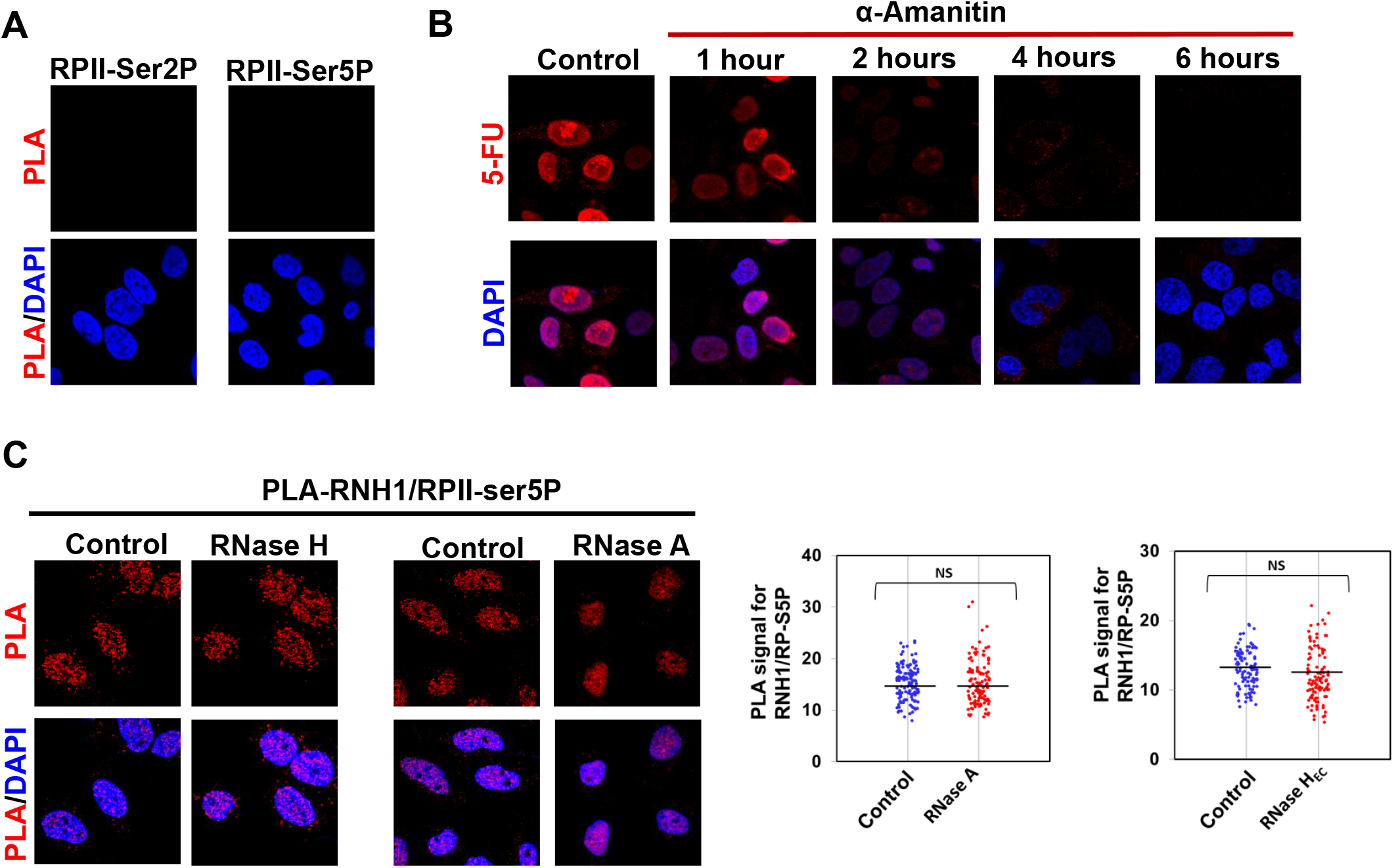
A. Single antibody controls of RPII-ser5P and RPII-ser2P used as negative control for PLA in Fig.2C. B. 5’-FU labeling (ant-BrdU immunostaining) in Hela cells upon treatment with α-Amanitin, a transcription inhibitor, for indicated times. α-Amanitin is known to bind to RNAP II with very high affinity near catalytic active site and inhibits nucleotide incorporation and translocation of transcript (Bushnell et al.,2002). Treatment of Hela cells with α-Amanitin (50 μg/mL), a transcription inhibitor, lead to significant down regulation of nascent RNA within 1-2 hours as indicated by FU labeling. Incubation of 2 hours was used for PLA reactions in Fig. 2C. C. PLA showing colocalization of RNH1 with RPII-ser5P (RP-S5P) after treatment of permeabilized growing Hela cells with RNase A or *E. coli* RNase H_EC_.

**Fig. S3.**
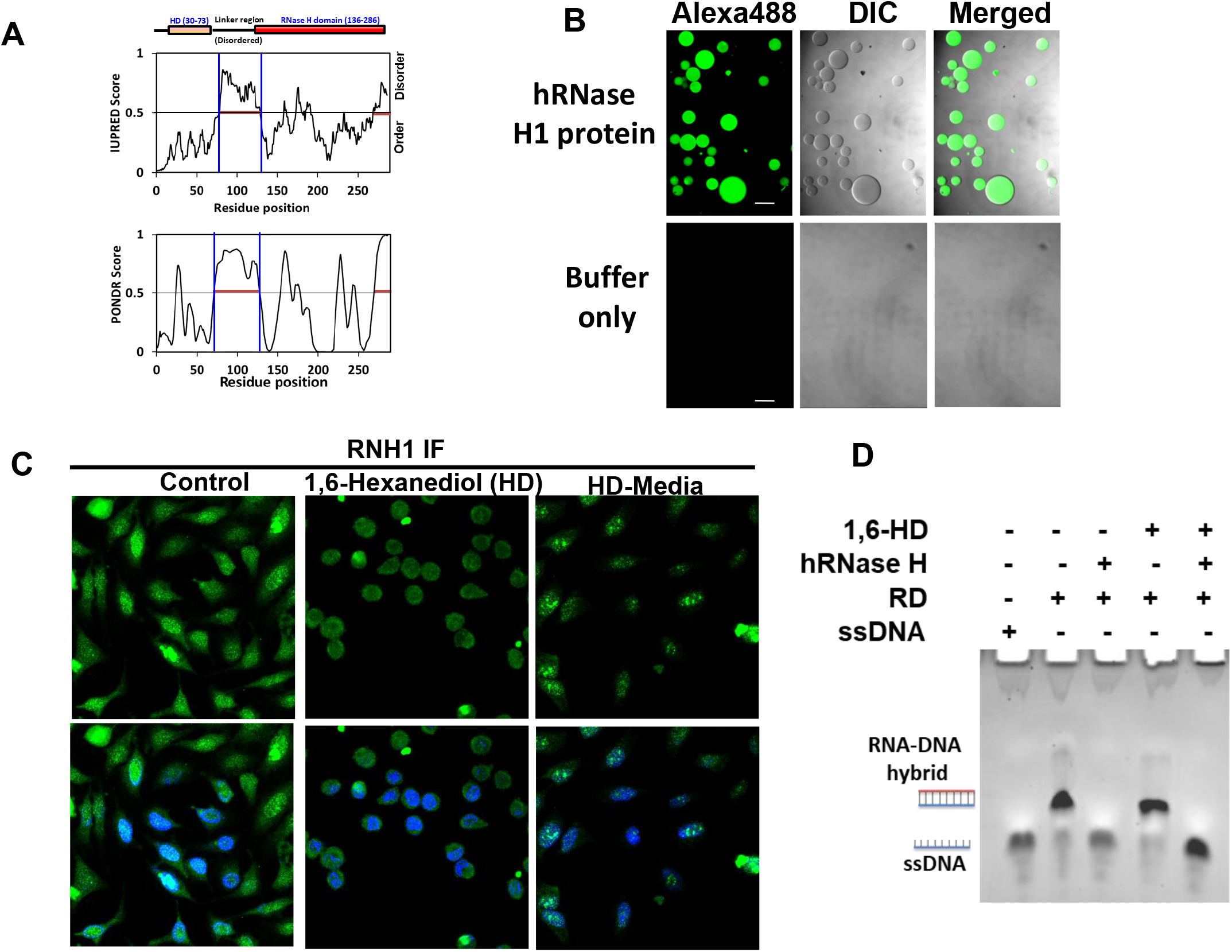
A. PONDR (Xue et al., 2015) and IUPRED2A (Mészaros et al., 2018) Plots aligned to different domains of RNH1. B. DIC and fluorescent confocal images showing Sypro labelled purified RNH1 protein and buffer only; Scale bar (100 µm) is shown by white line. C. RNH1 immuno-staining in untreated (control), 1,6-hexanediol (HD) treated, HD followed by media addition (HD+M). D. Effect of HD treatment on catalytic activity of purified human RNase H1.

**Movie S1**: A movie showing bleaching and following recovery of the fluorescence (Alexa488) during FRAP assay of Sypro labelled purified human RNH1 protein

**Table S1.**
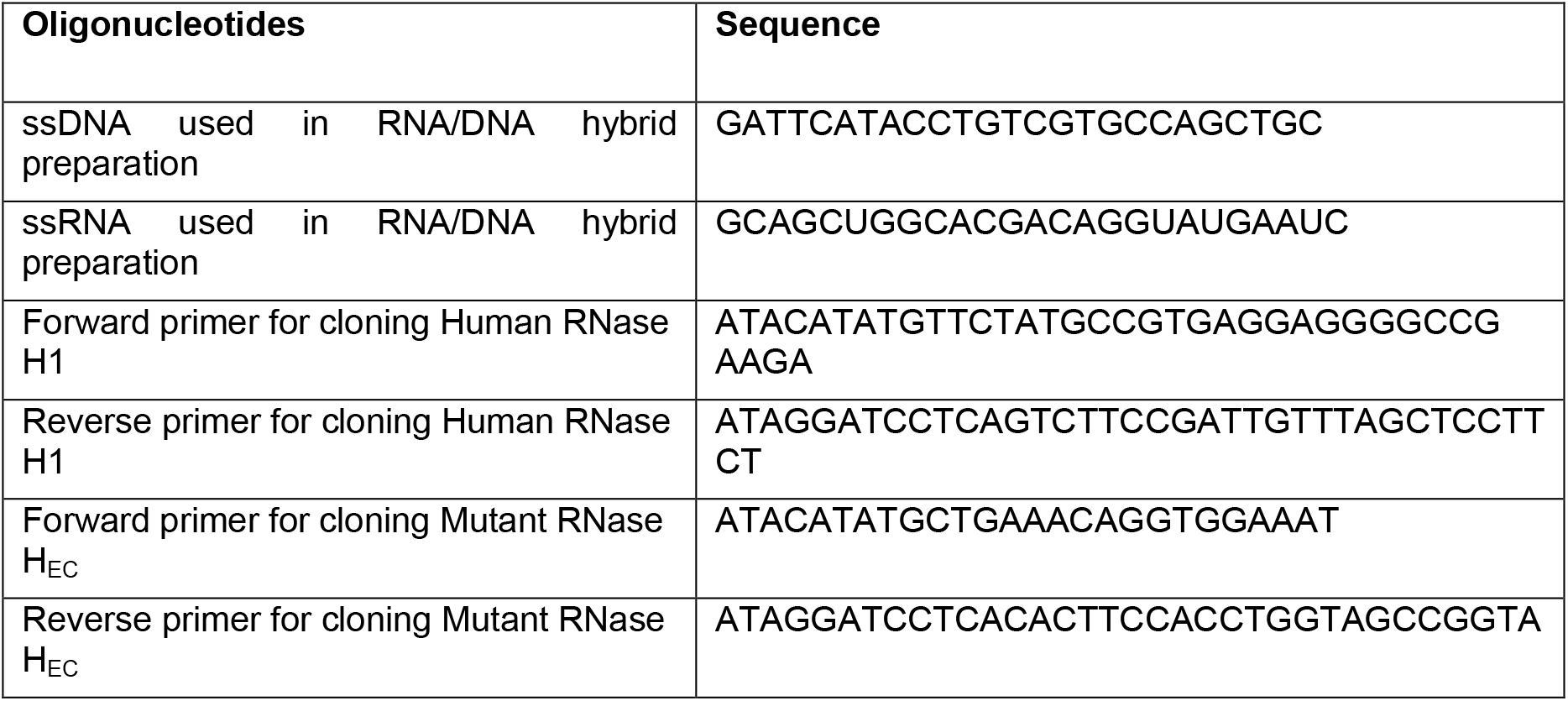
List of oligonucleotides used in this study

